# GABARAP Like-1 enrichment on membranes: Direct observation of trans-homo-oligomerization between membranes and curvature-dependent partitioning into membrane tubules

**DOI:** 10.1101/348730

**Authors:** Isabelle Motta, Nathan Nguyen, Helene Gardavot, Diana Richerson, Frederic Pincet, Thomas J. Melia

## Abstract

The Atg8/LC3/GABARAP protein family has been implicated in membrane remodeling events on the growing autophagosome. In particular, each of these proteins can form a protein-lipid conjugate that has been shown *in vitro* to drive liposome aggregation and in some cases membrane fusion. Furthermore, yeast Atg8 has been described as a curvature sensing protein, through its natural capacity to concentrate on highly curved membranes. A key advance with yeast Atg8, was the introduction of Giant Unilamellar Vesicles (GUVs) as an *in vitro* support that could allow membrane deformation and tethering to be observed by simple microscopy. Further, micromanipulation of an individual GUV could be used to create local areas of curvature to follow Atg8 partitioning. Here, we use a recently developed method to decorate GUVs with the mammalian Atg8 protein GABARAPL1 and establish the generality of the observations made on yeast Atg8. Then we apply double micromanipulation, the capture and positioning of two independently prepared GUVs, to test elements of the mechanism, speed and reversibility of mammalian Atg8 protein-mediated tethering. We find that the membranes adhere through GABARAPL1/GABARAPL1 homotypic trans-interactions. On a single membrane with two regions with significantly different curvatures we observed that the regions of higher curvature can be enriched up to 10 times in GABARAPL1 compared to the planar regions. This approach has the potential to allow the formation and study of specific topographically-controlled interfaces involving Atg8-proteins and their targets on apposing membranes.

## Introduction

Macro-autophagy, hereafter referred to simply as autophagy, is an intracellular degradative pathway in which cytoplasm or specific cytoplasmic cargos are sequestered in a growing cup-shaped double-membrane organelle called the isolation membrane (IM) or phagophore ^1^. This structure eventually closes upon itself, forming the mature organelle known as the autophagosome and sealing the cargos within. The autophagosome then delivers this cargo to the lysosome for degradation ^2^. How the isolation membrane elongates to envelope cargos and how it eventually closes upon itself in a membrane fission reaction are each unanswered mechanistic questions, but the major protein players involved have been described. In particular, the ubiquitin-like Atg8/LC3/GABARAP protein family has long been closely associated with both events.

Soluble Atg8/LC3/GABARAP proteins are recruited to the IM and become anchored through a covalent bond to the lipid, phosphatidylethanolamine (PE) ^3 4, 5^. When lipidation is prevented by disruption of critical upstream genes or expression of dominant negative proteins, most forms of autophagy are dramatically impaired while IM structures accumulate ^6–9^. Furthermore, when Atg8-PE/LC3-PE numbers are reduced in the cytoplasm of yeast and various mammalian cell lines, the resulting autophagosomes are smaller and a larger proportion of intermediates remain open ^10–12^,^13^, suggesting that Atg8-PE/LC3-PE is important in terminal membrane dynamics events like membrane elongation or pore closure. However, neither the lipidation event nor the mammalian Atg8 family is absolutely essential to the completion of autophagy ^14–16^; when the family is knocked out or lipidation is blocked, the autophagosomes that still form are delivered into the lysosome, but the efficiency and speed of closure and lysosome delivery are reduced (reviewed here ^17^). This suggests that these proteins do not play a central mechanistic role in autophagosome closure (for example these proteins are unlikely to be machines driving membrane fission or fusion ^17, 18^) but instead are an auxiliary facilitator of membrane dynamics. The mechanism of their facilitation is still being established, but in other systems, facilitators often include proteins that scaffold the fusion machinery or participate in membrane tethering.

In a paradigm-setting study, Ohsumi and colleagues reconstituted the lipidation of Atg8 onto small PE-rich liposomes and discovered that Atg8-PE can function in simple systems as a membrane tether^11^. This result has been reproduced by several groups^18, 19^ and extended to the mammalian and C. elegans homologs of Atg8, LC3 and GABARAP ^18–21^. An intriguing possibility is that membrane tethering could be used either to recruit vesicles to the growing IM (to facilitate membrane growth) or to initiate closure of the autophagosome by pulling the edges of the dilating rim together. These two potential tethering events present different physico-chemical membrane properties; closing a dilating rim requires tethering molecules to accumulate at the curved surface of the cup-like structure and would represent the close apposition of essentially homogenous surfaces (i.e. each side of the rim presents the same lipid and protein compositions); in contrast, tethering a vesicle has no inherent demand for curvature-recognition, and is inherently a heterogeneous interface, with the IM and incoming vesicle having different lipid compositions and possibly different Atg8-PE populations. Indeed, whether Atg8-PE is even resident on an incoming vesicle is unknown. Furthermore, because the size of each autophagosome rather than the number of total autophagosomes is reduced with lower Atg8-PE/LC3-PE formation, it is likely that the surface density of lipidated Atg8 proteins is integral to the process of membrane expansion, but how and whether surface density contributes to membrane tethering is unknown.

Here, we reconstitute the human Atg8 homolog, GABARAPL1, onto giant unilamellar vesicles (GUVs) to resolve individual tethering interfaces and then use this system to investigate whether *cis* or *trans* assemblies of these proteins ordinarily form, whether *trans* assemblies are necessary to support tethering, and whether the local concentration of these proteins is influenced by membrane structural perturbations consistent with strong curvature.

## Results and discussion

### GABARAP-dependent tethering of small liposomes

In humans, the Atg8-like family includes multiple LC3 proteins (LC3A, LC3B, and LC3C) and GABARAP proteins (GABARAP, GABARAPL1, GABARAPL2). Each member of the Atg8 family can be covalently attached to PE via ubiquitin-like Atg7-and Atg3-dependent reactions in cells and animals ^22–26^. To study the role of the Atg8 family independent of other cellular factors, the lipidation of Atg8 family proteins has also been reconstituted onto model membranes including small liposomes through the same ubiquitin-like reaction (e.g. ^11, 19, 21, 27–30^, and Figure 1A). Light scattering measurements and phase microscopy show that formation of Atg8-PE and LC3-PE is correlated with a quick and robust aggregation of the liposomes, establishing a natural capacity in these proteins for membrane tethering ^11,18, 20, 21^. Atg8-PE-mediated tethering was also observed on much larger and molecularly flat giant unilamellar vesicles (GUVs) ^31^. How Atg8-PE and LC3-PE drive membrane tethering is uncertain.

**Fig. 1.**
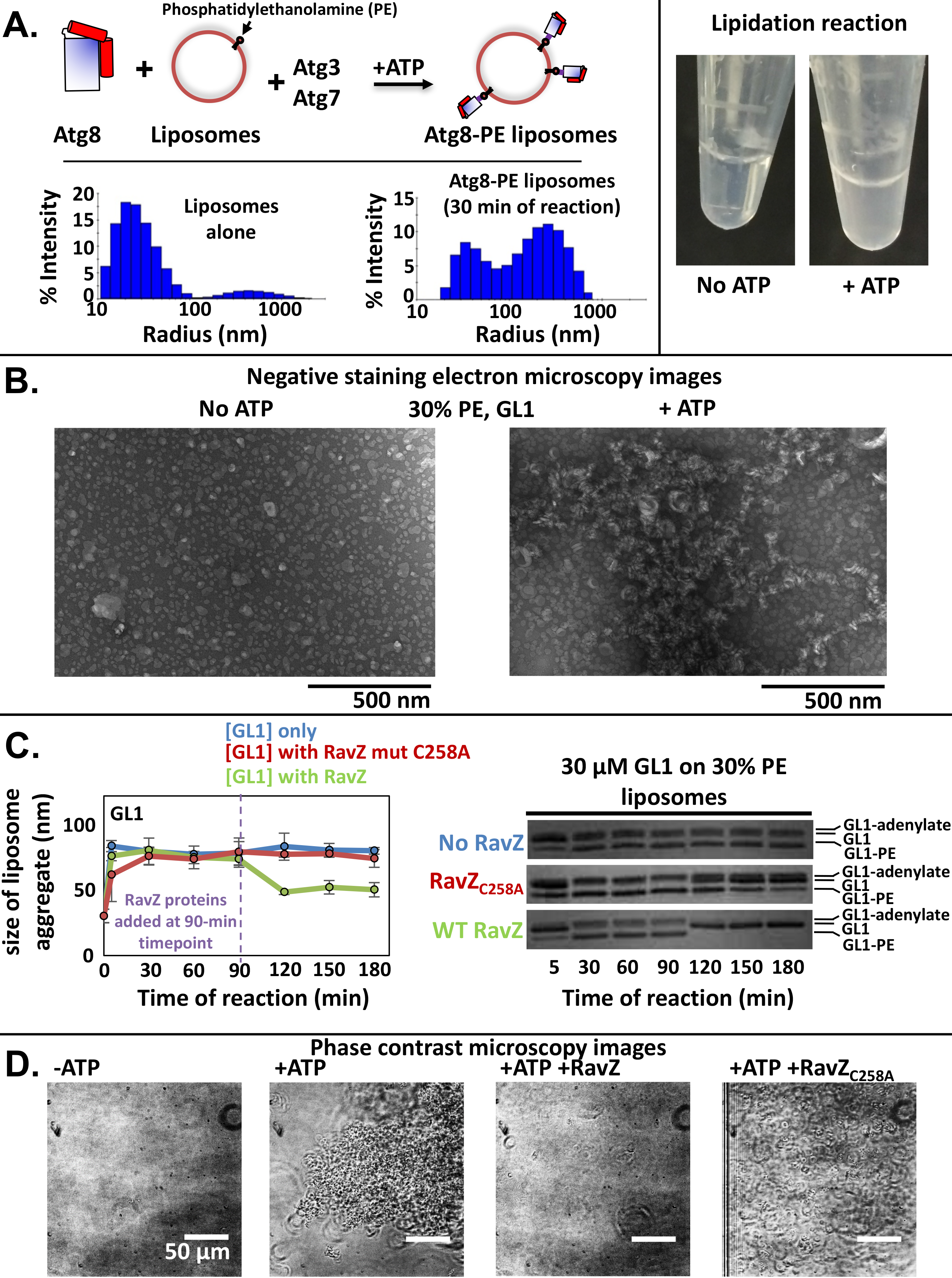
In vitro lipidation of GL1 proteins drives membrane tethering. (A) 30% PE or 55% PE small unilamellar vesicles (SUVs) were mixed with mammalian Atg8-lipidation machinery, GABARAPL1 (GL1) and ATP at 37°C for 90 minutes. GL1-dependent tethering is apparent as an increased cloudiness in the sample (right). Quantitative measurement of membrane aggregation is done via Dynamic Light Scattering (left). As the reaction progresses, the distribution of radius readings of liposomes shifts to larger sizes. (B) Transmission electron microscopy reveals dramatic membrane aggregation of liposomes lipidated with GL1 proteins after 90 minutes, in which aggregates range from hundreds of nanometers to microns in size. (C) To test whether membrane tethering by GL1-PE is reversible, the protease RavZ and a catalytically impaired mutant were introduced to the reactions. RavZ-dependent cleavage (gels on right) corresponds to a drop in DLS radii measurements back to pre-lipidation levels. (D) Imaging with phase contrast microscopy further confirms the ATP-dependent formation of vesicle aggregates and the RavZ-dependent reversibility of this GL1-PE controlled membrane tethering.

To study tethering in more detail, we first reconstituted the lipidation of three GABARAP family members onto membranes of different size and composition. Lipidation of GABARAP (GR), GABARAPL1 (GL1) and GABARAP L2 (GL2) is robust and essentially complete in our reactions by 30 minutes (Sup. Fig 1). At 37 degrees C and on very small liposomes produced by sonication, the lipidation efficiency is essentially the same for liposomes with high (55%) or moderate (30%) phosphatidylethanolamine surface densities, consistent with previous results ^28^.

To follow lipidation-dependent tethering, we used a number of different assays, each illustrated in figure 1 with GL1. As liposomes interact, they create large clusters that scatter light much more effectively. At high liposome concentrations, this can be easily observed with the naked eye in microcentrifuge tubes (Fig. 1A, right). To follow the time course of tethering, we use Dynamic Light Scattering (DLS) which can reveal the size of individual liposome aggregrates at early time points and under dilute liposome conditions (Fig. 1A, left). In a reaction mixture that includes liposomes, GL1, ATG3 and ATG7 but no ATP, DLS describes a narrow distribution of liposome sizes around a peak of about 35 nm radius, consistent with previous estimates of sonicated liposomes measured in this way. Upon addition of ATP to drive GL1-PE formation, these clusters grow to several hundred nanometers in diameter indicating the gross accumulation of at least tens of liposomes into clusters. These large structures are also obvious in negative stain electron microscopy (Fig. 1B); without ATP, liposomes are basically round and individual structures but after ATP addition, individual liposomes resemble flattened cisternae making large contact areas with 2 or 3 adjacent liposomes assembled into large aggregates.

At low liposome dilutions, DLS measurements suggest that GL1-PE drives a relatively homogeneous formation of small clusters with a radius of between 50 and 100 nm (Fig. 1C blue line, Fig. 2), consistent with only a few liposomes remaining in close apposition (but note that even under these conditions, some material is still in very large aggregates). The apparent homogeneity of most of the sample provides a convenient system to test other aspects of tethering. First, we note that liposome aggregates do not continue to grow after lipidation is complete (compare Fig. 2 and Sup. Fig. 1). Thus, under these conditions, GL1-PE is used essentially as fast as it is produced to drive tethering. There is also no apparent tethering when lipidation is blocked, even at very high GL1 concentrations (Sup. Fig. 2) suggesting GL1 cannot bridge the interface unless it is covalently attached to one membrane. Second, although lipidation is only modestly sensitive to PE composition on these small liposomes (Sup. Fig. 1), tethering is more dramatically impacted such that the higher PE leads to larger aggregates. This would be consistent with a portion of GL1-PE directly engaging the apposing bilayer via a membrane-embedding motif as was first demonstrated by Elazar and colleagues ^20^. However, this is also consistent with the possibility that 55% PE membranes support additional membrane dynamics including membrane fusion ^18, 19^. Indeed, on small extruded liposomes, we see that 55% PE supports fusion for GL1, GL2, and LC3 while 30% PE does not (Sup. Fig. 3).

**Fig. 2.**
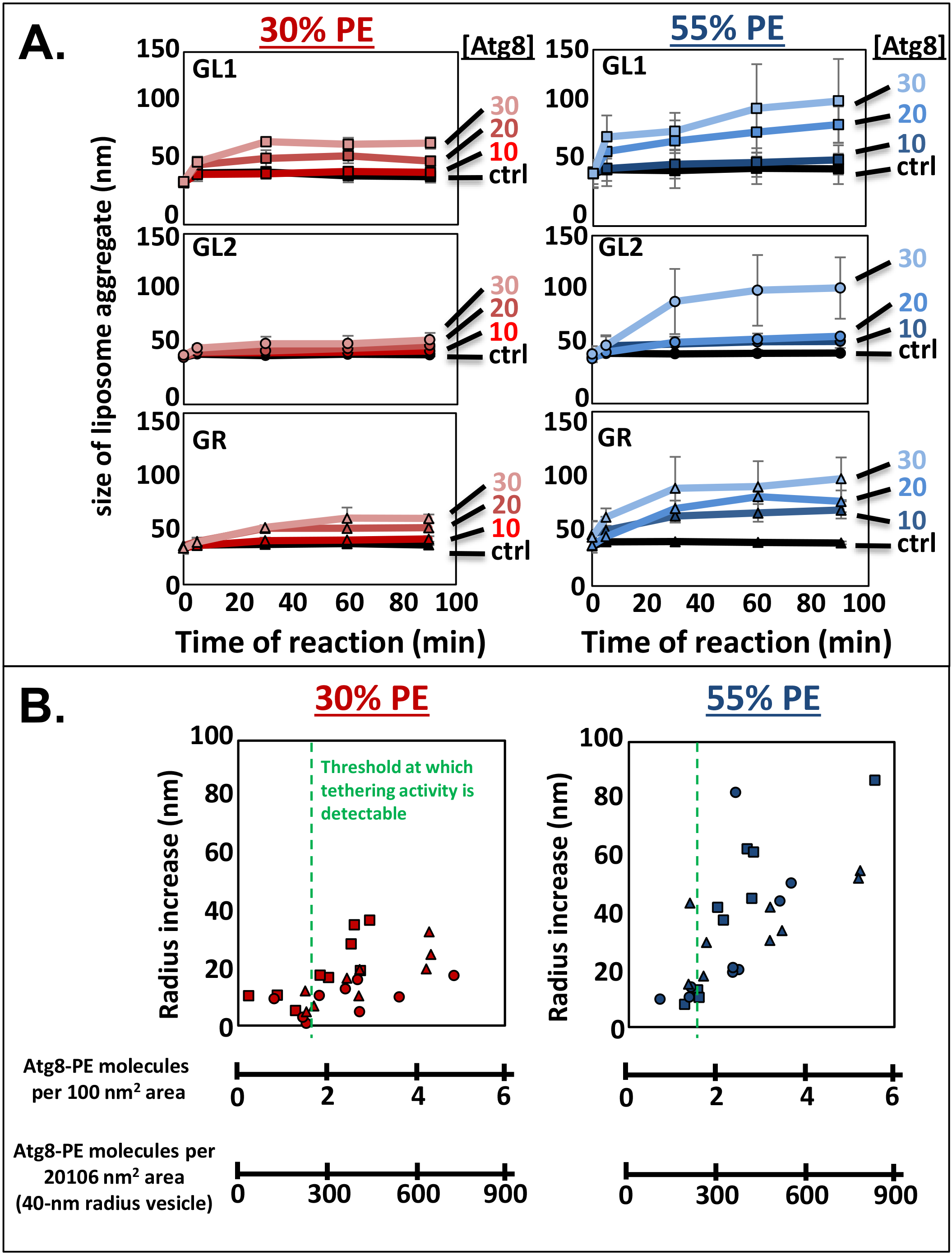
Membrane tethering is promoted by higher surface densities of GABARAP family proteins. (A) Lipidation reactions were set-up as in Fig. 1A and run at 37°C. At the indicated times, samples were pulled off and diluted and immediately subjected to DLS. The average distribution values for membrane radii increased with time and with GR/GL1/GL2 concentrations. Tethering was more efficient at higher membrane PE compositions, conditions which also promoted more efficient lipidation (Sup. Fig. 1). (B) To assess the minimum critical threshold of GABARAP family proteins in which tethering activity becomes apparent, the percentage of GR-PE vs. total GR in the system was quantified by densitometry, and the GR-PE concentration was compared to the total lipid concentration in the experiment to yield GR-PE molecules/100 nm^2^ area. For convenience, we also plot the number of GR-PE molecules that would be present at this density on a 40-nm radius liposome which approximates biologically-relevant structures like COP-II vesicles. This surface density of GR-PE was then plotted against the increase in vesicle radius detected by DLS.

Finally, we tested whether the GL1-PE dependent interface is accessible to outside modifiers and even potentially reversible. RavZ is an anti-autophagy protease that specifically targets the lipidated form of GABARAP and LC3 family members, cleaving the peptide bond between the lipid-anchored glycine and the aromatic amino acid before it ^32^. We drove lipidation and tethering of 30% PE liposomes for 90 minutes at 37 degrees before adding RavZ to the sample. From SDS-PAGE gels of the reactions, we could not detect any remaining GL1-PE in the RavZ treated sample at the next time point we measured (120 minutes; Fig. 1C and Sup. Fig. 4), suggesting that all of the GL1-PE in the sample remains accessible to exogenous protein. Likewise, via DLS measurements the liposome aggregates are rapidly disassembled following RavZ treatment. At very high liposome concentrations, large aggregates are apparent by phase microscopy (Fig. 1D), but these are also rapidly and completely reversed to undetectable small structures upon RavZ addition.

The surface density of GABARAP family proteins can be varied simply by changing the amount of GABARAP included in the reaction; from 10-30 μM GABARAP family proteins, we observe progressively more formation of lipidated proteins and in each case the reaction is exhausted by about the 30 minute mark. Tethering becomes more efficient at high surface densities (Fig. 2). This increase in productivity could reflect either a role for high surface densities during the formation of an interface (e.g. because collisional impacts are more productive) or could reveal a requirement for many copies of GL-PE in maintaining the stability of the interface. To distinguish between these two concepts, we used an impaired form of RavZ (RavZ_C258A_) in which the catalytic cysteine is mutated to an alanine. This mutation reduces its enzymatic activity by several orders of magnitude and thus provides a tool to slowly deplete GL1-PE from already tethered liposomes. With this treatment we observe that even as the starting GL1-PE is reduced by 90% (Sup. Fig. 4), liposome tethers remain intact (Fig. 1C/D). This implies that the high surface densities required to support tethering in vitro (Fig. 2) likely reflect limitations in initiating tethering but once tethers are formed the high avidity provided by tens of copies of GL1-PE is sufficient to maintain a complex.

### GABARAPL1 tethering on Giant Unilamellar Vesicles (GUVs)

Our results suggest that a minimal amount of GL1-PE is able to maintain a tethering interface once adhesion has been established (Fig. 1C) and that this interface is relatively broad (Fig. 1B); the EM images are consistent with a large surface of the liposomes being in contact with one another. Indeed, we expect that if the protein has an affinity for the apposing membrane, it should concentrate at the interface and promote further adhesion in a feed-forward manner. To directly observe this interface, we next tested GL1-PE tethering of Giant Unilamellar Vesicles (GUVs) in an assay similar to previous studies of Atg8 ^29–31^. GUVs are large enough (10-100 μm) to directly visualize by light microscopy. To follow GL1-PE on the surface of the GUV, we mutated the amino-terminus of GL1 to include a cysteine (the only cysteine in the protein) which we can chemically modify to include a small Alexa-488 fluorophore (Fig. 3A). The fluorescently modified form of the protein remains competent for *in vitro* lipidation (Fig. 3B), and still exhibits the same concentration-dependence to both lipidation and tethering as we observe for the unlabeled form of the protein (compare Fig. 3C with Fig. 2).

**Fig. 3.**
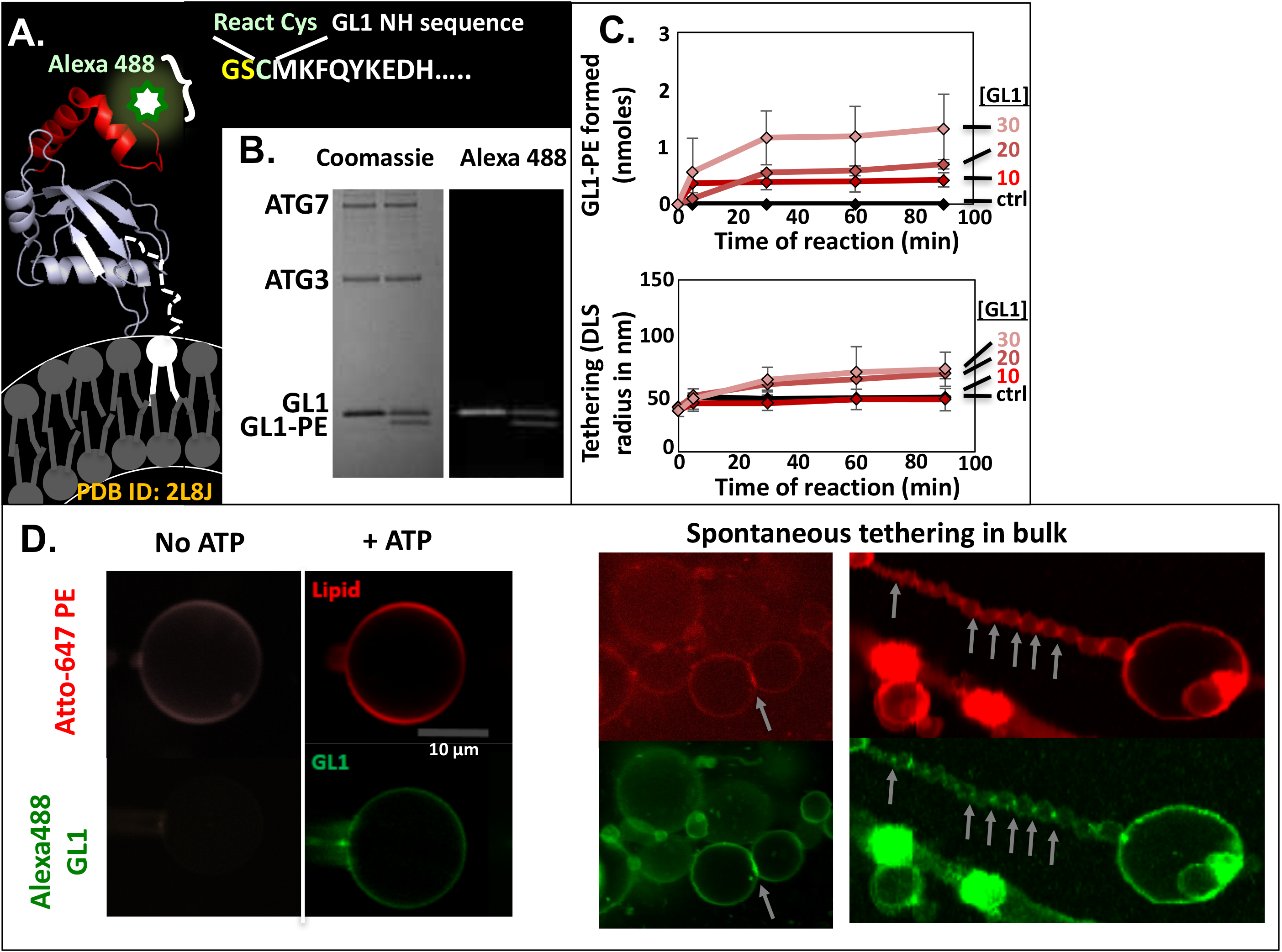
GABARAPL1-mediated tethering of Giant Unilamellar Vesicles (GUVs) (A) GL1 sequence was altered near the N-terminus to include a single cysteine residue which was then modified with the dye Alexa Fluor488 C5-maleimide via maleimide-thiol coupling (producing Alexa488-GL1). Cartoon model was produced from the PDB map of GL1 (ID 2L8J). (B) A lipidation reaction was run as in Fig. 1 on liposomes with 30 mol% PE. Both GL1 and GL1-PE molecules are visible by coomassie staining and Alexa 488 fluorescence. (C) Quantification of lipidation via densitometry and tethering activity via DLS of Alexa488-GL1 reactions (as in Fig. 2 and Sup. Fig 1). (D) GUVs were produced via the osmotic shock method with and without ATP to generate GUVs with and without incorporated Alexa488-GL1-PE, respectively. The osmotic shock method produces fields of GUVs (right) that can then be subjected to micromanipulation in order to isolate single GUVs (left). Arrows indicate regions of Alexa488-GL1-PE concentration at sites of GUV-GUV contact.

As the autophagosome lipid composition is still unknown we considered GL1 behavior in highly fluid membranes composed entirely of the unsaturated glycerophospholipids: dioleylphosphatidylcholine (DOPC at 69 mol%), and dioleylphosphatidylethanolamine (DOPE at 30 mol%). We also include a trace population of fluorescent lipid to visualize the bilayer (DOPE-ATTO647N at 1 mol%). *In vitro* enzymatic lipidation using only ATG3 and ATG7 is highly sensitive to membrane curvature or to the presence of local membrane instabilities created by very large surface densities of conical lipids ^28^, both properties that are inconsistent with GUV formation. Thus to facilitate GL1-PE integration into the essentially flat surface of a GUV, we recently described a novel GUV insertion protocol that relies upon first anchoring proteins in small liposomes and then fusing those vesicles via the osmotic-shock method ^33^. GL1-PE reconstituted on to GUVs with this approach remains highly mobile, and diffuses with fast kinetics similar to individual lipids ^33^, suggesting that Alexa488-GL1-PE exists as a monomer or potentially very small oligomer on GUVs.

Immediately after osmotic shock formation, the GUVs exist in a field of lipid structures closely associated with the bottom of the chamber but also mixed with large amounts of smaller liposomes in solution (Fig. 3D middle). The Alexa488-GL1-PE is integrated at a surface density of about 1 protein per 500 lipids and appears evenly distributed over the surface of each of the GUVs, except at the interface between large GUV structures where Alexa488 signal is increased (arrows). This increase is more obvious when the sample is washed to remove extraneous small vesicles (Fig. 3D right). At virtually every GUV-GUV interface, the accumulation of Alexa488 is obvious.

Importantly, when ATP is left out of the original coupling reaction, the GL1-free small liposomes can still be used to produce GUVs but they do not contain any Alexa488 signal (Fig. 3D. ″No ATP″) and thus are excellent controls in our system.

### Trans-interaction of the GABARAP protein

There are two classic paradigms for membrane tethering in cell biology. On the one hand, proteins may form multimers spanning the space between apposed membranes. These intermembrane complexes (*trans* interactions) provide specificity to the tethering reaction, as they are only supported when interacting proteins are present on both membranes. Just such a trans-intermediate complex has been inferred from the organization of GR in three-dimensional crystals. GR has been crystallized in two conformations, termed ″closed″ and ″open″, which are distinguished by the large movement of two alpha helices at the amino terminus ^34^. The open conformation reveals an association with a neighboring GR. This interaction appears to be conserved in solution when the salt concentration is high enough, but biochemical support for this model of interaction at membrane interfaces is lacking. In contrast, proteins with general lipid-binding motifs, including strong electrostatic attraction for polar head groups or highly hydrophobic membrane embedding structures, may drive the association of their anchored membrane with any other lipidic structure. A membrane-active motif that fits this description and is essential for tethering has been described for multiple homologs in the GABARAP family^20^, suggesting that *trans*-interactions may not be required. Addressing these two separate models is challenging in a bulk system using small liposomes; the best evidence that Atg8 is topologically required on both membranes during tethering comes from electron microscopy analysis of ATG8-PE LUVs with gold particles conjugated to Atg8 where interactions were largely restricted to situations where both vesicles were Atg8-PE positive ^11^. In principle, the single vesicle detection afforded by GUV experimental set-ups could directly establish whether protein-protein interactions are necessary.

To test whether GL1-PE is necessary on both membranes as in a *trans* configuration, we have employed both Alexa488-GL1-PE decorated and GL1-PE free GUVs in a GUV micromanipulation assay ^35, 36^. This experimental design allows us to capture single GUVs of known protein composition and then force these GUVs into direct apposition with a second well-described GUV of potentially different protein or lipid composition. For example, in Figure 4A we considered what happens to GUV protein distributions at vesicle contact sites when either both vesicles contain Alexa488-GL1-PE (top) or when only one vesicle contains Alexa488-GL1-PE (bottom). In this experiment, we used two different fluorescent lipids to distinguish vesicles arising from independent osmotic shock reactions. At the beginning of the experiment, one vesicle of each type (red or purple lipids) is separately micromanipulated via independent micropipettes at fully separate locations within the microscope dish and then moved into a common field of view in the confocal microscope. At this point, both the lipids and the Alexa488 are evenly distributed around the GUVs. Contact is initiated by micromanipulating the two GUVs into close apposition and this contact was maintained for 1-10 minutes. During this forced apposition, enrichment of the Alexa488-GL1-PE signal can be observed over the contact area (Fig. 4A, region 1), but only when both GUVs are GL1 positive. Across the contact area, we detected a 80 ± 40% increase in fluorescence beyond what is expected for the presence of two membranes, but only when Alexa488-GL1-PE was present on both GUVs. No recruitment was observed when only one GUV was decorated with GL1-PE (−16 ± 26%) ((Fig. 4A, region 2) and Fig. 4B). This strongly suggests that the GL1-mediated tethering is dependent upon a trans-interaction between GL1 proteins on apposing bilayers.

**Figure 4:**
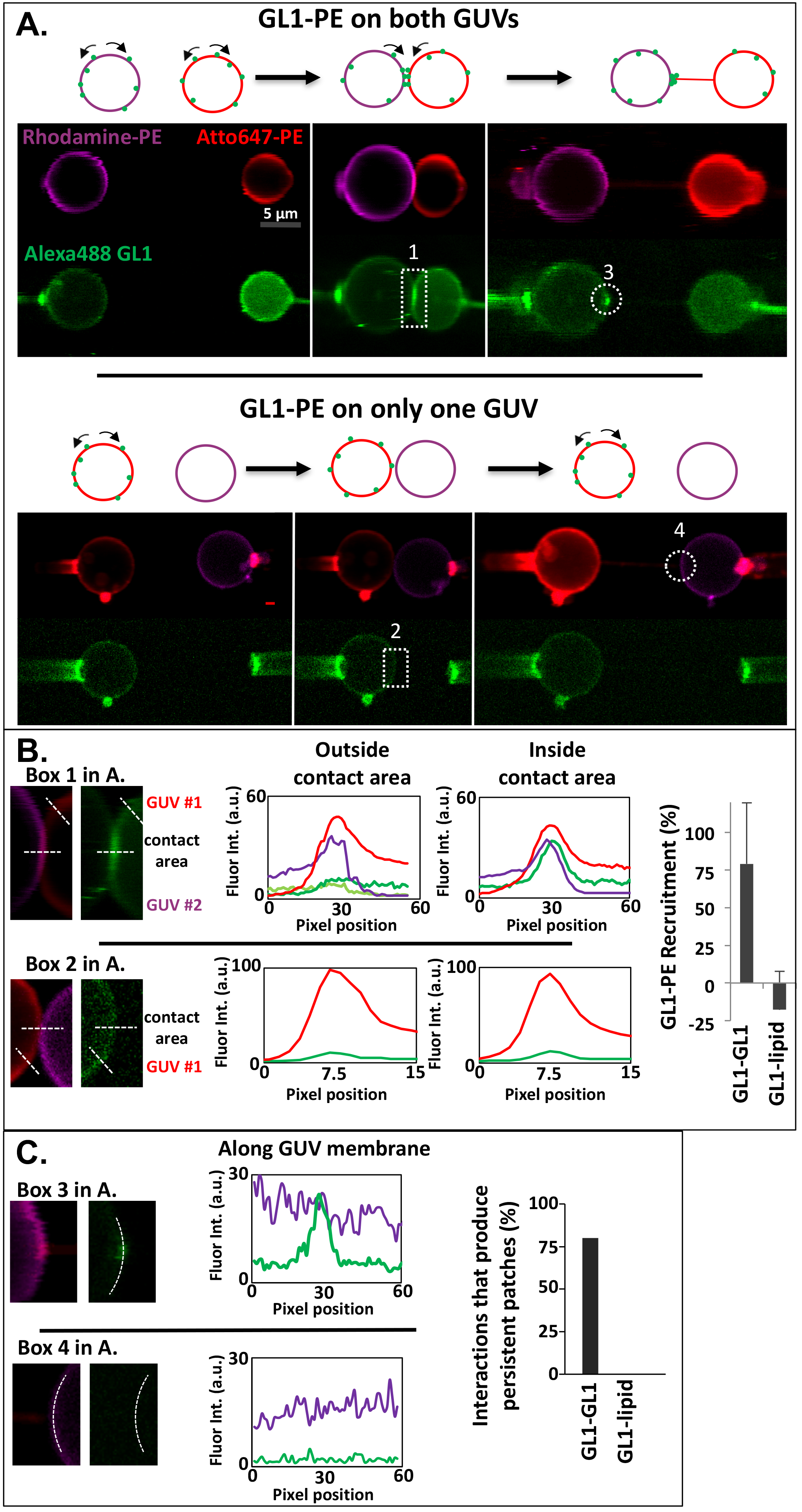
GABARAPL1-PE molecules on GUVs form trans-interactions. (**A**) Liposomes composed of 30% DOPE, 69% DOPC and 1% of either DOPE Atto-647 (Red) or DOPE-rhodamine (Magenta) were included in lipidation reactions with Alexa488-GL1 (Green) either without ATP (producing protein-free liposomes) or with ATP (to produce Alexa488-GL1-PE). These liposomes were subjected to the osmotic shock protocol ^33^. In brief, liposomes were dried and resuspended in water two times to form GUVs. In solution, two Alexa488-GL1-PE GUVs were micromanipulated together to make contact (1). Upon separation, a lipid tube formed between the two GUVs with a concentrated area of Alexa488-GABARAPL1 at one end of the tube (3). This protein patch often included a persistent accumulation of GL1-PE signal as expected for a GL1-PE dependent tethering mechanism. Scale bars, 5 μm. (**B**) The micromanipulation assays make use of both GL1-PE decorated and GL1-PE free GUVs. GUVs were micromanipulated into direct apposition and contact was maintained up to 10 minutes. Line scans across individual membranes or across contact sites revealed the average fluorescence intensity at these locations. Quantification of fluorescence intensity detected an 80% increase in Alexa-488 fluorescence at contact sites beyond what is expected from the simple summing of two decorated membranes when GL1-PE was present on both GUVs. No recruitment was observed when only one GUV was decorated with GL1-PE. (**C**) Quantification of fluorescence intensity across the protein patch suggests a further concentration of GL1-PE at these sites. These persistent accumulations of a GL1-PE into an adhering protein patch occurred ~80% of the time when both GUVs were decorated with GL1-PE and never when one GL1-PE decorated GUV was apposed to one GL1-PE free GUV.

After adherence, we next tested whether the tethering is reversible by forcibly separating the GUVs. Surprisingly, following separation a tube often formed from one of the interacting GUVs leading to a small area of persistent association between the two structures. This site of persistent association usually also resulted in a very strong accumulation of Alexa488-GL1 (Fig. 4A, region 3 and Fig. 4C), but again only if GL1-PE was present on both membranes (Fig. 4C). Thus, the adherence provided by GL1-PE is stronger than the limits of membrane deformation and one GUV will deform (producing a tube) to maintain the interaction. Furthermore, the site of contact appears to shrink even as the fluorescence intensity increases. This persistent contact thus is very likely approaching the limits of GL1-PE surface density that can be supported by the membrane.

Despite the strong adherence we observe, we were not able to detect any exchange of the two fluorescent lipids between the apposed GUVs in this system, again suggesting that GL1-PE does not support membrane fusion. Notably, an inability to fuse GUVs was also described by Knorr *et al.* with Atg8-PE ^31^, implying that both lipid composition and some membrane curvature are likely necessary to support the Atg8-mediated fusion observed on small liposomes.

### No cis-interaction of GABARAPL1-PE and YFP-GABARAPL1 on flat membranes

Whether the formation of a ″tethering-active″ GL-PE population first requires oligomerization in *cis* of multiple GL1-PE proteins on each individual membrane is unclear. Atg8 has long been known to form multimers as a consequence of lipidation, and mutants that disrupt this organization impair tethering and fusion^11^. In addition, the stark puncta that we observe from GUV-GUV separation resembles the kinds of protein aggregates that sometimes form in *cis* on membranes and could suggest a collapse to a stable low-affinity cis-oligomer. In our GUV system, Fluorescence Recovery After Photobleaching (FRAP) experiments established that GL1 is freely diffusible and unlikely to form large cis-structures ^33^, however FRAP is a bulk measurement, and we could miss small oligomers like dimers or trimers if they are present as part of a mix of different oligomeric states. In order to determine the range of stable available GL1-PE oligomeric states that normally form on a single membrane, we next performed single-oligomer bleaching experiments using a Total Internal Reflection Fluorescence (TIRF) microscope of Alexa488-GL1-PE incorporated into lipid bilayers formed directly on glass coverslip supports (Sup. Fig. 5A). In this experimental design, Alexa488-GL1-PE liposomes are first formed and then added to a dried lipid film in order to massively reduce the protein:lipid ratio. The final Alexa488-GL1-PE: lipid will be less than 1:10^9^. Then liposomes are spread directly onto clean glass coverslips and spontaneously fuse to form a large bilayer amenable to TIRF analysis. The GL1-PE surface density should be low enough such that each GL1-PE assembly will appear as a single fluorescent dot which will represent either individual proteins or strongly associating GL1-PE oligomers. During observation, the fluorescent spots undergo quantal bleaching steps with each step representing loss of one fluorophore (indicative of one protein) since the probability that two fluorophores would bleach within the same time frame is low. The analysis of this step-bleaching reveals that 87% of the Alexa488 spots contain only a single Alexa488 dye while about 13% contain two molecules (Sup. Fig. 5B). Because our labeling efficiency of Alexa488 to GL1 is only about 50%, this result suggests that ~75% GL1-PE are monomers and ~25% in dimers, but are unlikely to form stable oligomers of larger sizes. Importantly, GUVs and supported bilayers are morphologically comparable (i.e. both are essentially flat bilayer assemblies) and reveal similar behaviors of GL1-PE despite being generated by dramatically distinct experimental approaches (hydration shocking versus glass-dependent liposome spreading) and thus it is unlikely that in-plane oligomerization is an intrinsic property of the GL1-PE protein.

### Recruitment of the protein into highly-curved sub-domains of lipid bilayers

If GL1-PE mediated tethering is used in closing the autophagosome, the protein will have to reside on the highly curved rim of the organelle. The lipidation of Atg8/LC3 on liposomes is strongly sensitive to membrane curvature ^28^, and several other autophagy complexes recognize or depend on local curvature (reviewed here ^37^), suggesting proteins in this pathway may have natural affinities for strongly deformed membranes. Indeed, Atg8-PE has been shown to prefer highly curved surfaces ^31^. Our micromanipulation system is ideal for establishing whether GL1-PE also preferentially distributes into these types of membranes. To produce regions of high curvature, we pulled tubes of defined and very small radii from our otherwise essentially flat GUVs, using streptavidin coated beads that bind to DSPE-biotin present in the GUV bilayer (0.5 mol %) (Fig. 5A). The radius of the pulled tube can be controlled by careful changes in the aspiration of the micromanipulation pipette holding the GUV. By measuring both the changes in the length of the tube and the length of the membrane extension (″tongue″) within the micropipette during aspiration these tube radii can be precisely determined ^38^ (Fig. 5A). Over the samples considered in our experiments, the radius of the tubes varies from 15 to 37 nm. This range is convenient as it explores the curvature landscape of most small biological objects as vesicles including the highly curved rim of the developing autophagosome. Once the tube is pulled, we can determine the recruitment of the Alexa488-GL1-PE into the tube relative to the concentration of fluorescent lipids in the tube which exhibit no curvature-dependent local enrichment (Fig. 5B). We observed a small consistent enrichment of Alexa488-GL1-PE into our tubes. For ten independent samples, enrichment was detected in each case with an average increase of about 5-fold over the surface density of Alexa488-GL1-PE in the GUV itself. We could not detect a further curvature dependence across the narrow range of radii we tested, indicating that while the difference between a flat GUV surface and the tube is dramatic, our set-up cannot resolve meaningful differences between 14 and 40 nm radii. This range of recruitment is similar in magnitude to that detected by Knorr and colleagues and suggests a subtle curvature sensitivity is a conserved property of the Atg8 family.

**Fig. 5.**
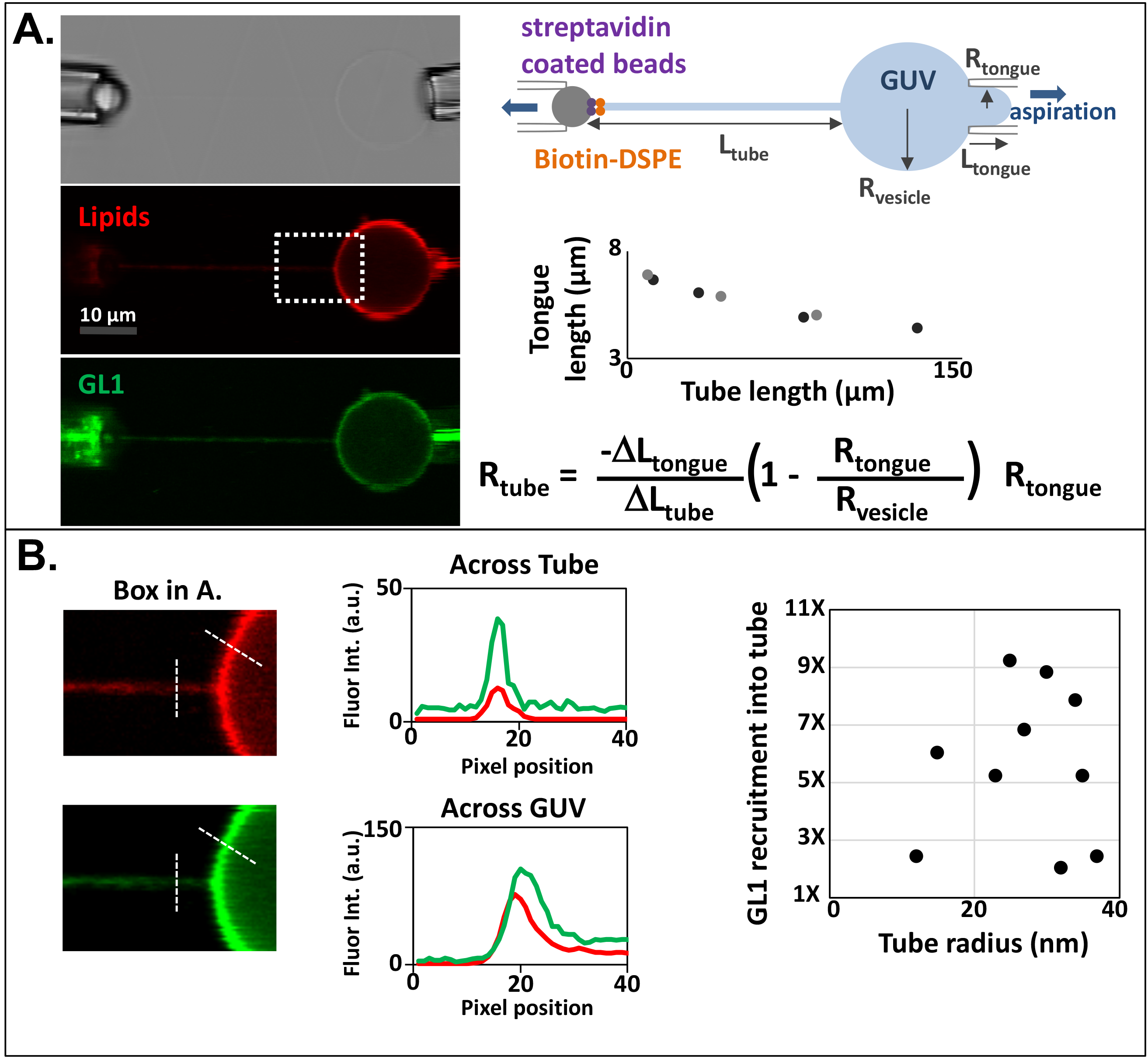
GL1-PE molecules are recruited onto highly curved membranes. (**A**) To investigate whether GL1-PE is naturally recruited to membranes that are highly curved, tubes of defined and very small radii (between 15 and 35 nm) were pulled from GUVs using streptavidin coated beads that bind to DSPE-biotin present in the GUV bilayer (0.5 mol %). The curvature of the tube is the inverse of its radius, and the radius of the pulled tube can be controlled by careful changes in the aspiration of the micromanipulation pipette holding the GUV. To calculate the tube radius, both the changes in the length of the tube and the length of the membrane ″tongue″ within the micropipette were measured. Scale bar: 10 μm. (**B**) To calculate the recruitment of GL1-PE in the tubes, fluorescence intensities were measured. Once the tube is pulled, the measurement was made between recruitment of the GL1-PE into the tube relative to the concentration of fluorescent lipids in the tube which exhibit no curvature-dependent local enrichment. From these collected results, GL1-PE molecules are recruited 5.5x more often in tubes, compared to flat surfaces. However, there is no clear correlation between GL1 recruitment as a function of membrane curvature over the narrow range of curvatures tested here.

## Conclusion

The autophagic cup is a structurally asymmetric organelle that necessitates equally asymmetric distributions of local biochemistry. For example, membrane growth must be coordinated with organelle closure at or near the leading edge of the cup surface, while cargo capture must be promoted or stabilized on the interior face of the cup. How these asymmetries are determined remains unknown but at the heart of each of these activities is the lipid-anchored form of one or more homologues of Atg8. Here we explore whether GABARAPL1 alone or in combination with particular membrane cues, is inherently capable of defining local asymmetries in protein distributions. Our results strongly support a model where GABARAPL1 accumulates at sites of membrane-membrane apposition, provided that GABARAPL1 is anchored to PE on both of the interacting membranes. The effect of this accumulation is to create a tight seal between the two apposing membranes that remains intact even when the vesicles are forcibly separated. Despite this strong interaction, we do not observe any evidence of lipid-mixing in this system, thus how and whether such accumulations would be resolved in vivo is uncertain.

One appealing hypothesis, built upon early ideas from the Ohsumi lab, is that GABARAPL1 could bridge some portion of the open fission pore to facilitate organelle closure. If so, the completion of the fission reaction would render the apposing membranes as a single structure, eliminating the opportunity for trans-assemblies and possibly dispersing the accumulated GABARAP proteins. Our previous results suggest that if such a mechanism were driven solely by Atg8, LC3 or GABARAP proteins, it would require a highly unstable or highly energetic substrate membrane, structures which are generally not favored in cells due to their natural fusogenicity. However, one would expect the very highly curved nature of the autophagosome rim to provide this basal instability. Importantly, we also demonstrate an inherent affinity of GABARAPL1-PE for highly curved membrane surfaces, using tubular membranes as a structural proxy for the autophagosome rim. This result is in good agreement with a recent study on Atg8-PE^31^, suggesting that accumulation at curved surfaces may be a general property of the broader Atg8 protein family.

LC3 and GABARAP proteins engage a large number of cytosolic factors more efficiently and perhaps specifically only when LC3 and GABARAP are lipidated ^39^. Thus, some property of membrane association changes the protein reactivity of these autophagosome markers. It appears that the increased surface density of these proteins may be sufficient to specifically capture cytosolic proteins harboring many low affinity LC3/Atg8 interaction sites ^40^, and possibly to alter the activity of membrane targeted enzymes ^41^, and thus local concentrations of these proteins are clearly important. Whether actual oligomers are forming however has been uncertain. Our results demonstrate that at low surface densities, GABARAPL1 is predominantly monomeric and even at higher densities, we do not observe protein diffusion rates comparable to dramatically higher order assemblies, and thus we would suggest that GABARAPL1 is unlikely to intrinsically enter into cis-homo-oligomers. How other proteins might facilitate such an oligomerization, consistent with the recruitment and stabilization of Atg8 by Atg5-Atg12/Atg16 remains to be determined.

## Acknowledgements

This work was supported by grants from the NIH (GM1000930 and NS063973; TJM), a Burroughs Welcome Fund travel grant to facilitate the international collaboration (BWF ID #1014056), the pre-doctoral program in Cellular and Molecular Biology (T32GM007223; supporting NN), and a doctoral fellowship (FRM FDT20140931036; IM).

## Materials and Methods

### Materials

Phospholipids: 1,2-dioleoyl-sn-glycero-3-phosphocholine (DOPC), 1,2-dioleoyl-sn-glycero-3-phosphoethanolamine (DOPE), 1,2-dioleoyl-sn-glycero-3-phosphoethanolamine-N-(lissamine rhodamine B ph) (DOPE-rhodamine), 1,2-dioctadecanoyl-sn-glycero-3-phosphoethanolamine-biotin (DSPE-biotin), 1-palmitoyl-2-oleoyl-sn-glycero-3-phosphocholine (POPC), and L-α-phosphatidylinositol from bovine liver (bi-PI), were purchased from Avanti Polar Lipids. ATTO647-dioleoyl-phosphoethanolamine (DOPE-ATTO647) were purchased from ATTO-Tec.

Making liposomes: Chloroform (BD), Borosilicate Glass 10×75mm (Fisher Scientific), polycarbonate membranes (Whatman), LipSoFast-Basic extruder (Avestin), PrecisionGlide™ Needle 1.2mm × 40mm (BD). Virsonic 600 (VirTis) microtip sonicator.

Dye: Alexa Fluor488 C5-maleimide (Alexa488) was purchased from Life Technologies.

GUV specific items: Dialysis cassettes Slide-A-Lyzer™ MINI Dialysis Device, 7K MWCO, 0.1 mL (Thermo Fisher). 35-mm dish with 200-mm glass bottom (MatTek).

Lipidation items: NuPAGE LDS Sample Buffer 4x (Thermo Fisher), 12% bis-Tris SDS-PAGE gels (Novex), NuPAGE^®^ MES SDS Running Buffer 20x (Thermo Fisher), SeeBlue Plus2 Pre-stained Protein Standard (Thermo Fisher), Coomassie Blue Stain Imperial Protein Stain (Thermo Scientific). XCell SureLock™ Mini-Cell Electrophoresis System (Thermo Fisher). BioDOT Universal Fit Pipette Tips (Dotscientific). Hoefer EPS 2A200 Power Supply C.

Protein expression items: BL21-Gold (DE3) Competent Cells (Agilent Technologies), Luria Bertani broth (LB) media and plate, carbenicillin antibiotics, Isopropyl β-D-1-thiogalactopyranoside (IPTG), Ethylenediaminetetraacetic acid (EDTA)-free protease inhibitor cocktail tablets, thrombin, precision protease, glutathione-agarose beads (Sigma)

Others: Tris (American Bioanalytical), Trizma hydrochloride (Tris HCl) (American Bioanalytical), Sodium chloride (American Bioanalytical), Glycerol (J.T. Baker), TCEP (Sigma-Aldrich). Magnesium chloride 6-hydrate, crystal (J.T. Baker), Calcium chloride, Bovine serum albumin (BSA) (Thermo Fisher), Nycodenz, Dithiothreitol (DTT) (American Bioanalytical), Adenosine-5′-trisphosphate, disodium (American Bioanalytical). 5×41 mm Ultra-Clear™ ultracentrifuge tubes (Beckman Coulter)

sw55Ti rotor and Optima™ L-90K Ultracentrifuge (Beckman Coulter). F14 rotor and Sorvall Evolution RC Centrifuge (Thermo Fisher).

### Protein expression & purification for human Atg8 homologs and ATG3

Human GABARAPL1/GABARAPL2/GABARAP (mammalian Atg8 homologs) were cloned in the PGEX-2T GST vector and mouse ATG3 was cloned into PGEX-6p vector. The mammalian Atg8 homologs are ordinarily expressed in a pro-form that requires processing from Atg4 to expose the C-terminal glycine in order for lipidation to take place. To facilitate lipidation in our in vitro system, GABARAPL1/GABARAPL2/GABARAP are expressed with COOH-terminal truncations such that the reactive glycine is fully exposed and ready for interaction with ATG7 and ATG3. The mammalian Atg8 homologs and ATG3 proteins were expressed in BL21-Gold (DE3) Competent Cells (Agilent Technologies). Cells were cultured in 4L Luria Bertani Broth (LB) media with 1:1000 carbenicillin (50 mg/mL) and induced with 0.5 mM final concentration IPTG. Bacterial pellets were treated with EDTA-free protease inhibitor cocktail tablets in either thrombin buffer (20 mM Tris pH 7.5, 100 mM NaCl, 5 mM MgCl2, 2 mM CaCl2. 1 mM DTT) for Atg8 proteins or precision protease buffer (50 mM Tris pH 7.5, 150 mM NaCl, 1 mM EDTA, 1 mM DTT) for ATG3. Cells were broken in aa cell disrupter and the lysate was incubated with glutathione beads for 3 hours at 4°C. Beads were washed several times and then incubated with Atg8 cutting buffer (10 uL thrombin + 500 uL thrombin buffer + 0.5 uL DTT + 500 uL beads) or ATG3 cutting buffer (25 uL precision protease + 500 uL precision protease buffer + 0.5 uL DTT + 500 uL beads) to cut the proteins from GST tags overnight. Purified proteins were stored in 20% glycerol at −80°C.

### Protein expression & purification for GABARAPL1 5′ cysteine

Visualization of GABARAPL1 on Giant Unilamellar Vesicles (GUVs) is possible via specific dye labelling at a unique cysteine amino acid near the amino terminus. GABARAPL1 was mutated with Quik Change II Site-Directed Mutagenesis Kit (Agilent Technologies) to include the cysteine immediately before the start methionine of the natural GABARAPL1 sequence in the pGEX-2T GST vector. In this organization, there remains two additional amino acids N-terminal to the cysteine which derive from the thrombin cleavage site used to release GST. Expression and purification of this GABARAPL1 5′-cysteine protein was similar as described above with the Atg8 mammalian homologs, except instead of 1 mM DTT we used 0.2 mM TCEP in the buffer as TCEP is necessary to protect the cysteine for the dye labelling reaction.

### Protein expression & purification for human ATG7

Human ATG7 in pFastBac vector was from Sloan-Kettering (kind gift of X. Jiang). The plasmid was transformed into Bacmid DNA and PCR was used to check for the bacmid that obtained the insert. 2 ug DNA was used to infect 8×10^5^ SF9 cells by using Cellfectin II two times to increase viral titer to 5×10^6^ pfu/ml. 1×10^8^ pfu/mL of SF9 cells were infected with virus and they grew for 72 hours. Cells were treated with EDTA-free protease inhibitor cocktail tablets in the lysis buffer (20 mM Tris pH 8, 500 mM NaCl, 20 mM Imidazole, 1 mM DTT, 10% glycerol), sonicated with the Virsonic 600 (VirTis) microtip on for 3 minutes (30 sec on, 30 sec off. Speed 3.5) and centrifuged at 18000 rpm for 1 hour. The lysate was incubated with 1 mL Nickel resin (Ni-NTA Agarose) for 2 hours at 4°C. The beads were washed with the wash buffer (20 mM Tris pH 8, 300 mM NaCl, 20 mM Imidazole, 1 mM DTT) three times and eluted with the elution buffer (20 mM Tris pH 7.5, 300 mM NaCl, 500 mM Imidazole, 1 mM DTT). Purified proteins were stored in 20% glycerol at −80°C.

### Protein expression & purification for *Legionella pneumophila* RavZ

Recombinant *L. pneumophila* RavZ constructs were expressed in BL21-Gold (DE3) Competent Cells (Agilent Technologies). Cells were cultured in 4L Luria Bertani Broth (LB) media with 1:1000 carbecillin (50 mg/mL) and induced with 0.2 mM final concentration IPTG, then the temperature was lowered to 18°C and cells were harvested 20 hours later. Bacterial pellets were treated with EDTA-free protease inhibitor cocktail tablets in precision protease buffer (50 mM Tris pH 7.5, 150 mM NaCl, 1 mM EDTA, 1 mM DTT). Cells were smashed and proteins were bound with glutathione beads for 3 hours at 4°C. Beads were incubated with the cutting buffer (25 uL precision protease + 500 uL precision protease buffer + 0.5 uL DTT + 500 uL beads) to cut the proteins from GST tags overnight. Purified proteins were stored in 20% glycerol at −80°C.

### Dye labeling for GABARAPL1 5′ cysteine

GABARAPL1 was labeled with Alexa Fluor488 C5-maleimide (Life Technologies) through the amino terminal cysteine. 20 uL of 500 μM GABARAPL1 was diluted in 80 uL Tris NaCl buffer (50 mM Tris HCl, 100 mM NaCl) and 11 uL of 60 mM TCEP then mixed. After 5 min incubation at room temperature, 8 mM fluorescent dye was dissolved in 10 uL DMSO. The mixture was protected from the light and slowly mixed at room temperature for 2 hours. The labeled GABARAPL1 was then dialyzed with dialysis buffer pH = 7.6 (100 mM NaCl, 50 mM Trizma hydrochloride) overnight at 4°C to remove the excess free dye.

### Production of Small Unilamellar Vesicles (SUVs) for Dynamic Light Scattering (DLS) and imaging experiments

30 mol% DOPE, 60% POPC, and 10% bi-PI or 55 mol% DOPE, 35% POPC, and 10% bi-PI were mixed together. The mixture is dried with N^2^ and vacuumed for 1 hour to form a lipid film. The film was resuspended in SNH buffer pH 7.6 (20 mM Tris, 100 mM NaCl, 5 mM MgCl_2_) to make 10 mM final concentration. The liposomes underwent 7x freeze-thaw cycles with liquid nitrogen and 37°C water bath to form unilamellar liposomes. Liposomes were extruded 21 times through polycarbonate membranes (Whatman) with 50-nm pore size using the LipSoFast-Basic extruder (Avestin) at room temperature. Liposomes were sonicated with the Virsonic 600 (VirTis) microtip on ice for 10 minutes (1 min each. 1 sec on, 1 sec off. 50% power). Liposomes were stored on ice for future usage.

### *In-vitro* reconstituted lipidation reactions to decorate GABARAPL1-PE on SUVs

*In vitro* lipidation reactions were performed as previously described ^1–3^. Each reaction is 100 uL in volume and contains purified ATG3 (2 μM), ATG7 (2 μM), GABARAPL1 (10-30 μM) or its homologs, mix with ~20 to 40-nm radius liposomes (sonicated SUVs) (2 mM lipid composed of 30 mol% DOPE, 60% POPC, and 10% bi-PI or 55 mol% DOPE, 35% POPC, and 10% bi-PI), 1 mM dithiothreitol, and 1 mM ATP. Reactions were run at 37°C for 90 minutes in SNH buffer (20 mM Tris at pH 8, 100 mM NaCl, and 5 mM MgCl_2_).

### DLS of SUV proteoliposome tethering

Each sample in the experiment was prepared as described above to make GABARAPL1-PE (and its homologs) decorated SUVs. The *in vitro* lipidation reactions proceed in 37°C water bath for 90 minutes in SNH buffer. Time-points for the DLS readings were at 5, 30, 60, and 90 minutes. For each time-point, 5 μL of the reaction was mixed with 55 μL SNH buffer then subjected to laser at wavelength 600 nm at 25°C to acquire 10 radius readings. Membrane radii were measured using the Dyna Pro Titan Dynamic Light Scattering instrument (Wyatt Technology Group) and analyzed using the Dynamics V7 software (ver. 7.1.7.16).

### Gel visualization and quantification

15 uL samples were mixed with 4 uL 4x SDS-PAGE and boiled at 90°C for 5 minutes. Electrophoreses of samples were performed in 12% bis-Tris SDS-PAGE gels (Novex) running in 1x MES SDS Running Buffer (NuPage Invitrogen) for 55 minutes at 200V using the Hoefer EPS 2A200 Power Supply C. Samples were visualized with Coomassie Blue stain as per the manufacturer′s instruction (Imperial Protein Stain, Thermo Scientific). Band intensities of samples were quantified with ImageJ software as described ^1–3^. Gels were imaged with the VersaDoc Imaging System (Bio Rad) and analysed with the software Quantity One (vers. 4.6.9) - Protein Gels Coomassie Blue setting. Images were saved in JPEG format and analysed with ImageJ software. Gel analyses were done by making a rectangle encompassing the non-lipidated Atg8 and lipidated Atg8 bands. Plots were generated from this to show the density of contents of the rectangle over each band. Lipidation efficiency was calculated by taking the area of the Atg8-PE peak to divide over the area of the (Atg8-PE peak + Atg8 peak) and multiply by 100%.

### Visualization of SUV samples with transmission electron microscopy (TEM)

1 uL sample from the lipidation reaction described previously for the DLS experiment was diluted with 20 uL SNH buffer. The sample was deposited on a glow discharged carbon coated copper grid (Electron Microscopy Sciences) for 3 minutes, then incubated with 5 uL 2% (w/v) uranium formate stain two times for 1 minute each. Images were acquired with the JEOL USA JEM-1400Plus Transmission Electron Microscope with the 4k×3k CCD camera (Advanced Microscopy Technologies) at 80 keV acceleration voltage. Images analysis was done with AMT software.

### DLS and Phase Contrast Microscopy of RavZ delipidation and detethering

*In vitro* lipidation reactions were performed as previously described ^1–3^ for the DLS experiments. Samples were then incubated with RavZ at 90 minutes and were analyzed with DLS as described previously at 120,150, and 180 minutes. 20 uL of the sample was imaged with the Nikon Eclipse Ti-E inverted Microscope equipped with the spinning disk confocal camera Yokogawa CSU-W1 CFI and 60x oil objective. Image analysis was done with NIS-Elements and ImageJ software.

### Production of Large Unilamellar Vesicles (LUVs) for Giant Unilamellar Vesicles (GUV) imaging and micromanipulation experiments

30% DOPE, 1% DOPE Atto647 or DOPE Rho, 69% DOPC were mixed together. For tube pulling experiments, 0.5% DSPE-biotin was added to the composition (68.5% DOPC + 0.5% DSPE-biotin). The mixture is dried with N^2^ and vacuumed for 1 hour to form a lipid film. This film was resuspended in SNH buffer pH 7.6 (20 mM Tris, 100 mM NaCl, 5 mM MgCl_2_) to make 10 mM final concentration. The liposomes underwent 5x freeze-thaw cycles with liquid nitrogen and 37°C water bath to form unilamellar liposomes. Liposomes were sonicated with the Virsonic 600 (VirTis) microtip on ice for 5 minutes (30 sec on, 30 sec off. Speed 3). Liposomes were stored on ice for future usage.

### *In-vitro* reconstituted lipidation reactions to decorate GABARAPL1-PE on LUVs

*In vitro* lipidation reactions were performed as previously described ^18, 19, 28^. Each reaction is 150 uL in volume and contains purified Atg3 (2 μM), Atg7 (2 μM), Alexa488-GABARAPL1 (12-15 μM) or its homologs, mix with LUVs described above (2 mM lipid composed of 30% DOPE, 1% DOPE Atto647 or DOPE Rho, 69% DOPC or 68.5% DOPC + 0.5% DSPE-biotin), 1 mM dithiothreitol, and 1 mM ATP. Reactions were run at 37°C for 90 minutes in SNH buffer (20 mM Tris at pH 8, 100 mM NaCl, and 5 mM MgCl_2_).

Floation assay: Nycodenz liposome flotation was performed as described previously ^41^. Lipidation samples were mixed with 150 uL 80% Nycodenz (100% (w/v) Nycodenz in SNH buffer). Nycodenz gradients were established in 5×41 mm Ultra-Clear™ ultracentrifuge tubes (Beckman Coulter) with 300 μl of 40% reaction mixture at the bottom, followed by a 125 μl middle layer of 30% Nycodenz and a 40 μl top layer of 0% Nycodenz SNH buffer. Each tube was centrifuged at 48000 rpm in an sw55Ti rotor (Beckman) for 4 hours at 4°C. The liposomes with lipidated Alexa488-GABARAPL1 proteins were collected from the top portion of the gradient above the 30% Nycodenz mark.

### Hydration-Shocking Method for the production of GABARAPL1-PE decorated GUVs from LUVs

2 uL of GABARAPL1-PE containing LUVs were deposited on 35-mm dish (MatTek), dried under atmospheric pressure, rehydrated with 6 uL water, dried again, and hydrated a last time with 6 uL water to produce GABARAPL1-GUVs ^33^. The temperature for the hydration step should be carefully controlled, since the saturated lipid DPPC has a transition temperature equals to 41°C, the hydration should be done at a temperature ≥ 41°C; we set the temperature at 41°C to limit protein instability.

### Fluorescence microscopy of GUVs

GUVs were detected using bright field and confocal microscopy at room temperature. The experiments were carried out on a Leica SP5 Confocal Microscope equipped with a 10x and a 20x air objective, an Argon (488/543nm) and HeNe (633nm) lasers, and a 488/543/633 beam splitter. Image analysis was done with the LAS AF software.

### Micromanipulation and FRAP of GUVs

Micromanipulation was used to manuever independent GUVs and keep them static in solution. The internal diameter of the micropipettes was about ~5 μm and to prevent adhesion of the GUV on the glass, the pipette was incubated in 200 uL 10% BSA prior to use for 15 min. The aspiration to maintain the GUVs was controlled by hydrostatic pressure and kept to 20 Pa. Fluorescence recovering after photobleaching (FRAP) was performed on GUVs to monitor the diffusion coefficient of fluorescent lipid and fluorescent protein on the membrane. The fluorescence recovery curves were then analyzed with Mathematica 10.

---

**Supplementary Fig. 1. In vitro lipidation is sensitive to GABARAP family protein concentration and lipid composition.** (A). Representative examples of lipidation reactions. Each reaction contains 2 uM ATG3, 2 uM ATG7, 10, 20, or 30 uM GL1, GL2, or GR, 2 mM lipid (liposomes prepared by sonication and composed of 30 mol% DOPE, 60% POPC, and 10% bi-PI or 55 mol% DOPE, 35% POPC, and 10% bi-PI), 1 mM dithiothreitol, and 1 mM ATP. Reactions were run at 37°C for 90 min in SNH buffer (20 mM Tris at pH 8, 100 mM NaCl, and 5 mM MgCl_2_). Each reaction was run on a 12% SDS-PAGE gel and visualized by coomassie blue stain. The lipidated form of the protein migrates faster than the non-lipidated form on the gel. (B). Densitometry quantification of the lipidated form is determined in ImageJ and plotted as percentage of total GR. Each experiment was run in triplicatew error bars represent the standard deviation.

**Supplementary Fig. 2. High concentrations of soluble (non-lipidated) GR do not lead to tethering.** To test whether free GR can bind membranes and promote tethering, we compared GR at low (10 μM) and high (30 μM) concentrations in the presence of a lipidation enzymes but in the absence of ATP, against a moderate GR concentration (20 μM) in the presence of ATP. We used liposomes that were sonicated and contained 55% PE, thus maximizing the probability of curvature-dependent recruitment of ATG3 as well as any non-specific recruitment of other proteins including GR. The samples were run at 37°C for 90 minutes. Neither ATP-free condition lead to any evidence of tethering (A) or lipidation (B), while the ATP-containing reactions were effective at both.

**Supplementary Fig. 3. Mammalian Atg8 proteins only support membrane fusion on liposomes with high concentrations of PE.** (A). Cartoon of the fluorescence dequenching assay used to monitor lipid-mixing between liposomes. In brief, donor liposomes carry fluorescent lipids with both NBD and rhodamine fluorophores. Within the donor liposomes, the concentrations of these two fluorophores are high enough that the NBD fluorescence is naturally quenched by the rhodamine. Following lipid-mixing, the surface density of the rhodamine goes down, and NBD fluorescence can be detected. (B). Representative fusion experiment with GL1. 50 nm extruded liposomes were prepared with 10% bovine-liver phosphatidylinositol, either 30% or 55% DOPE, and completed with POPC. Donor liposomes also carried 1% NBD-PE and 1% rhodamine-PE. Liposomes were mixed at a 4:1 donor to target ratio. Lipidation reactions were then initiated by adding ATG3, ATG7, ATP and GL1 and run at 37°C for 90 minutes. Fluorescence was measured every 2 minutes. ″No ATG7″ samples are negative controls. (C). Summation of fusion experiments for three different mammalian Atg8 homologs. Experiments were conducted as in (B), using GL1, GL2 or LC3. Each experiment was run in triplicate and then the maximal fusion rate obtained in the experiment was plotted. Lines indicate average values at each point. Significant fusion was only detected in samples with 55% DOPE and containing a fully active lipidation mixture.

**Supplementary Fig. 4. GL1-PE in tethering interfaces remains accessible to GL1-targeted proteases.** Lipidation reactions were run at 37°C for 90 minutes with GL1 on 30% PE liposomes. At the 90-minute time point, samples were treated with the protease RavZ or the RavZ_C258A_ catalytic mutant. RavZ was able to cleave GL1. Representative gels are shown in Figure 1. Densitometry quantification of experiments done in triplicate are shown here with the amount of the lipidated form remaining after RavZ treatment plotted as percentage of total GL1.

**Supplemental Fig. 5. Single molecule bleaching confirms Alexa488-GL1-PE is not in large complexes.** (A) Cartoon of method: GL1 LUVs were prepared by lipidation as in figure 3. GL1 LUVs were then diluted with resuspension buffer and added to a film of dried lipids. The final GL1:lipid ratio in the sample was less than 1 GL1 per 10^9^ lipids. Liposomes were then spread on a clean glass surface to generate a bilayer. Provided GL1 is mobile, the very low density of protein to lipid ensures that individual GL1 or GL1 oligomers will appear as single fluorescent dots under the TIRF microscope (B). To test the oligomerization of GL1, we subjected each dot to bleaching over time. The dots undergo quantal bleaching steps consistent with the bleaching of one individual fluorophore/step. Across the population, the vast majority of spots correlated to a single fluorophore. Because GL1 labeling efficiency is only ~50%, this implies that each spot is likely either a monomer or dimer of GL1, but not a larger oligomer.

